# AlphaFold2’s training set powers its predictions of fold-switched conformations

**DOI:** 10.1101/2024.10.11.617857

**Authors:** Joseph W. Schafer, Lauren L. Porter

## Abstract

AlphaFold2 (AF2), a deep-learning based model that predicts protein structures from their amino acid sequences, has recently been used to predict multiple protein conformations. In some cases, AF2 has successfully predicted both dominant and alternative conformations of fold-switching proteins, which remodel their secondary and tertiary structures in response to cellular stimuli. Whether AF2 has learned enough protein folding principles to reliably predict alternative conformations outside of its training set is unclear. Here, we address this question by assessing whether CFold–an implementation of the AF2 network trained on a more limited subset of experimentally determined protein structures– predicts alternative conformations of eight fold switchers from six protein families. Previous work suggests that AF2 predicted these alternative conformations by memorizing them during training. Unlike AF2, CFold’s training set contains only one of these alternative conformations. Despite sampling 1300-4400 structures/protein with various sequence sampling techniques, CFold predicted only one alternative structure outside of its training set accurately and with high confidence while also generating experimentally inconsistent structures with higher confidence. Though these results indicate that AF2’s current success in predicting alternative conformations of fold switchers stems largely from its training data, results from a sequence pruning technique suggest developments that could lead to a more reliable generative model in the future.

## INTRODUCTION

AlphaFold2 (AF2) is a deep-learning-based algorithm that predicts a protein’s three-dimensional structure from its amino acid sequence, often with high accuracy (Jumper et al. 2021). AF2’s success at predicting one dominant conformation of an amino acid sequence has led to the development of algorithms for sampling conformational ensembles and alternative conformations with varying success (del Alamo et al. 2022; Jing et al. 2024; Kalakoti and Wallner 2024; Monteiro da Silva et al. 2024; Vani et al. 2023; Wayment-Steele et al. 2024; Zheng et al. 2024). Some of these approaches focus on predicting both conformations of fold-switching proteins, a class of globular proteins that remodel their secondary and/or tertiary structure in response to cellular stimuli (Porter and Looger 2018). Recent results indicate, however, that AlphaFold2 and AlphaFold3-based approaches are weak predictors of fold switching (Abramson et al. 2024; Chakravarty and Porter 2022; Chakravarty et al. 2024)

How AF2-based methods predict alternative protein conformations remains unclear. The abstruse nature of neural network-based predictions is a universal problem that has motivated approaches that render human-interpretable explanations for how deep learning approaches work (Lundberg and Lee 2017; Mehdi and Tiwary 2024; Ribeiro et al. 2016). Two such explanations for how AF2 predicts alternative conformations have been proposed: the Generative Explanation, which posits that AF2 has learned enough folding principles to consistently and accurately predict alternative conformations outside of its training set, and the Associative Explanation, which posits that AF2’s successful predictions of alternative conformations often depend on the structures it learned during training. In further detail, the Generative Explanation suggests that AF2 uses evolutionary couplings from its input multiple sequence alignment (MSA) to predict alternative conformations (Sala et al. 2023). In this framework, the information needed to specify a given fold is provided by the MSA, enabling AF2 to generate predictions of alternative protein conformations with high confidence regardless of whether they were in its training set. Alternatively, the Associative Explanation suggests that AF2 predicts alternative conformations from “memory” of structures learned during training (Bryant and Noe 2024), allowing it to associate related input sequences and/or MSAs with these structures (Chakravarty et al. 2024). These inputs do not necessarily provide enough information to enable AF2 to predict a new fold accurately, however (Schafer et al. 2024). Instead, structures learned during training foster accurate predictions from sparse sequence information, limiting AF2’s ability to reliably predict alternative conformations outside its training set (Chakravarty et al. 2024).

Here, we test how the AF2 network predicts alternative conformations of fold-switching proteins by running it with a new set of model weights trained on the dominant conformations of a subset of fold switchers but not their alternative conformations (**Figure 1, Supplementary Methods, Table S1**). Bryant and Noé trained this implementation of AF2 to predict alternative conformations and called it CFold (Bryant and Noe 2024). If the Generative Explanation is correct, we would expect CFold to predict these alternative conformations of fold switchers correctly and uniquely with high confidence, as AF2 does. If the Associative Explanation is correct, we would expect CFold to fail to predict accurate alternative conformations with high confidence and/or to predict incorrect structures with high confidence, indicated by per-residue local distance difference test plDDT scores ≥ 70 (or 0.7 for CFold). In all cases tested, we find that CFold fails to predict alternative conformations of fold-switching proteins outside of its training set reliably, supporting the Associative Explanation. However, predictions from a previously suggested sequence filtering technique (Schafer and Porter 2023) suggest developments that could lead to a more reliable generative model in the future.

**Figure 1.**
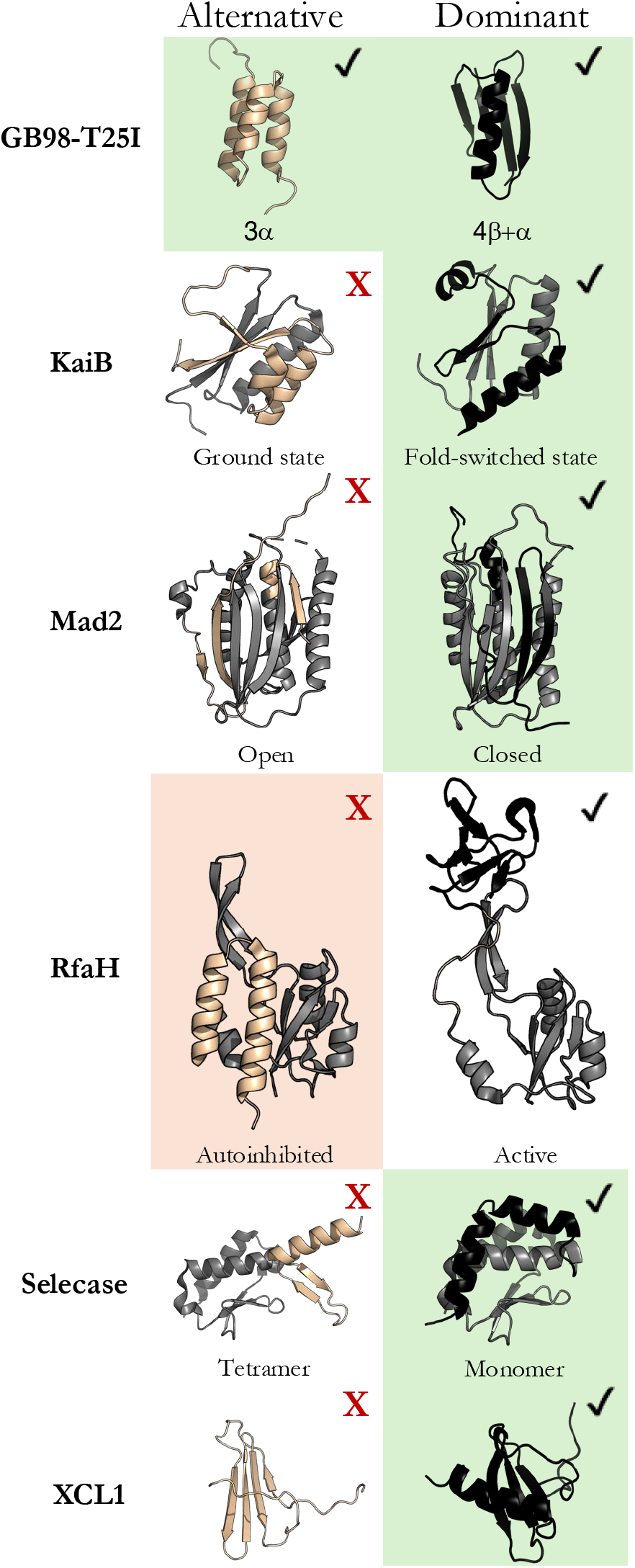
Alternative and dominant conformations of fold-switching proteins from 6 families. CFold predictions from full MSAs denoted with check boxes, those not predicted with red Xs; AlphaFold2 conformations predicted from full MSAs denoted with green and orange backgrounds. Green/red means AF2 and CFold predicted the same/different conformations. Fold-switching regions of proteins are beige and black.

## RESULTS AND DISCUSSION

CFold was tested on eight fold-switching proteins, six of which are from different fold families (**Figure 1**). Previous work suggested that AF2 memorized the alternative conformations of six of these during training and the dominant conformation of selecase; we added the eighth (GB98-T25I) because AF2 predicts both of its experimentally determined conformations accurately and with high confidence. GB98-T25I is an engineered fold-switching protein that reversibly transitions between a 3-α-helix bundle and 4β+α fold (He et al. 2012). Three experimentally characterized KaiB proteins, whose fold switching helps regulate cyanobacterial circadian rhythms (Chang et al. 2015), were also evaluated (*S. elongatus* KaiB, *R. sphaeroides* KaiB, *T. elongatus* KaiB). KaiBs belong to a protein superfamily (large clade of diverse homologous sequences) composed of both single folders and fold switchers. All its members are expected to fold into a dominant thioredoxin-like fold, but those that switch folds also assume an alternative ground state. The bacterial transcriptional regulator RfaH, whose C-terminal domain reversibly switches between a dominant β-sheet and an alternative α-hairpin, the eukaryotic mitotic spindle protein Mad2 (Mapelli et al. 2007), a bacterial Selecase that switches folds upon oligomerization (Lopez-Pelegrin et al. 2014), and the fold-switching chemokine XCL1 (Dishman et al. 2021) were also evaluated. It is important to note that “dominant” denotes the conformations that CFold predicts with highest confidence from deep MSAs, not the conformation that the apo protein assumes under native conditions. Previous work shows that these “dominant” conformations are predicted from rich coevolutionary information (Schafer and Porter 2023), explaining why CFold predicts them from full MSAs. A FoldSeek search (van Kempen et al. 2024) indicated that CFold’s training set contained all dominant conformations but only one alternative conformation of these fold-switching proteins (GB98-T25I, **Supplementary Methods, Table S1**).

Full-MSA sampling produced predictions of all structures in CFold’s training set: all dominant and the one alternative (**Figure 1**). Neither full-MSA sampling nor the MSA subsampling suggested to be used with CFold (Bryant and Noe 2024) produced any of the alternative conformations outside of CFold’s training set (**Figure S1**), despite sampling 800 structures/protein (**Table S2**). Notably, while AF2 predicts RfaH’s alternative helical conformation with high confidence from full MSA sampling, CFold consistently predicts its dominant β-sheet conformation with high confidence and fails to predict its helical conformation with either full-MSA sampling or suggested MSA subsampling (**Figure S1**). RfaH’s helical conformation is not in CFold’s training set, but it was likely in AF2’s (Chakravarty et al. 2024). These results support previous observations suggesting that AF2 memorized RfaH’s alternative helical conformation during training (Chakravarty et al. 2024).

Two other MSA-based approaches were used in an attempt to predict alternative conformations outside of CFold’s training set: sequence clustering based on similarity (Wayment-Steele et al. 2024) and subfamily filtering or pruning of the protein family tree (Schafer and Porter 2023). These approaches–used to generate an additional 1100-3500 structures–differ because: (1) sequence clusters are shallower, containing ≤11 sequences while filtered subfamilies contain dozens to hundreds of sequences and (2) sequence clusters do not necessarily preserve phylogenetic order while subfamily-filtered sequences do. Though AF2 successfully predicted the alternative conformations of some fold switchers using sequence clusters as input (Wayment-Steele et al. 2024), CFold did not accurately predict any alternative conformations with high confidence from them (**Figure S2, Supplementary Methods**). This result supports previous work indicating that the coevolutionary information from sequence clusters is insufficient to generate alternative conformations outside of AF2’s training set (Schafer et al. 2024).

Subfamily filtering enabled more predictions of conformations loosely resembling some alternative conformations, but it was sufficient to capture the alternative conformation of only one KaiB variant accurately and with high confidence (**Figure 2**). This indicates that evolution has likely selected for many of these alternative conformations (Schafer and Porter 2023), but the information provided by subfamily filtering is not sufficient to enable CFold to reliably predict alternative conformations of these fold switchers. For instance, CFold produced RfaH structures with alternative helical C-terminal domains (CTDs), though these predictions were low-confidence (plDDT < 0.6), and the structural quality was poor, with a best overall RMSD ≥ 12.0Å (**Figure S3A**). Furthermore, incorrect structures with mixed α-helix and β-sheet CTDs were produced with higher confidence than the correct α-helical form (**Figure S3A**). Thus, while subfamily filtering provides CFold with some information about RfaH’s alternative fold, it is not sufficient to produce an accurate structure, nor does it enable recognition of the alternative structure with high confidence. Further, CFold failed to accurately predict the alternative conformations of Mad2, Selecase, and XCL1 with high confidence (**Figure S3B**), while AF2 has been successfully achieved this for the alternative conformations of Mad2 and XCL1 (Chakravarty et al. 2024). It should be noted that our definition of a successful prediction is TM-score ≥ 0.6; if Bryant and Noé’s threshold of 0.8 is used then zero alternative structures were successfully predicted (**Figure 3**).

**Figure 2.**
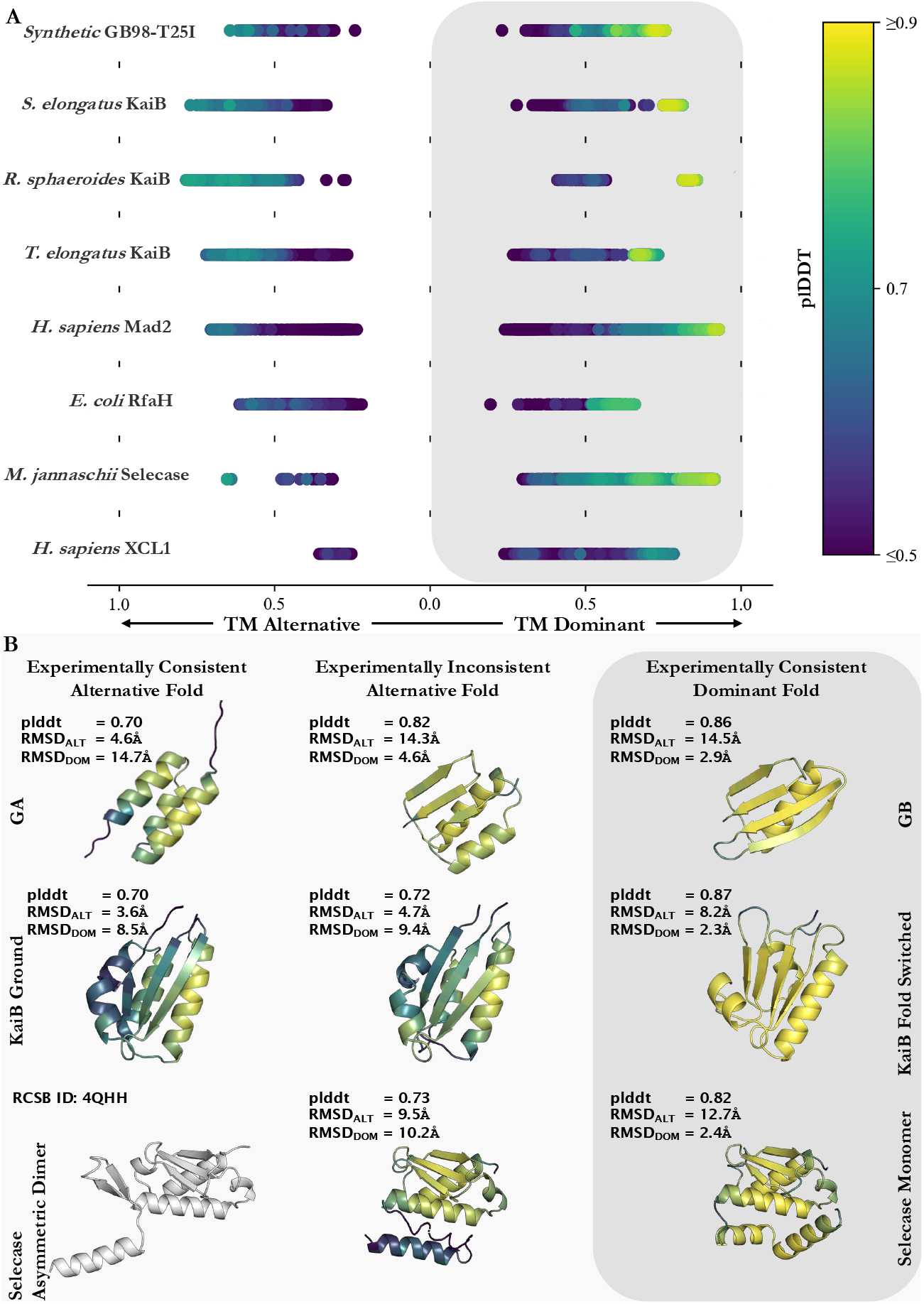
Pruning by sequence identity enables CFold to predict some experimentally consistent alternative structures with high confidence but also produces high-confidence structures inconsistent with experiment. (A) TMscores for ensembles of predicted structures created with CFold for fold-switching proteins. The predicted structure with the highest plDDT score defines the dominant predicted conformation for the CFold ensemble. Predicted structures within an ensemble are sorted by TMscore: if the TMscore is greater for the dominant conformation the value appears on the right side of panel A (gray box); otherwise, the value appears on the left-side of panel A. All TMscores are colored by the predicted structures average plDDT score. (* RfaH calculations are sorted by TMscore calculated for residues 118-155 to ensure conformations for this two-domain protein are organized by the fold-switching region. The TMscore plotted is the TMscore for the entire sequence compared to crystal structures. XCL1 calculations are on residues 1-65, the last 29 residues are unstructured and are not considered for TMscore or plddt score) Experimentally determined structures used for comparison in order from top to bottom (alternative/dominant) are 2LHC/2LHD, 2QKE_A/1T4Y, 4KSO_A/8FWJ_M, 2QKE_A/1T4Y, 3GMH_L/2VFX_L, 5OND_A/6C6S_D, 4QHH_A/4QHF_A, 2N54_A/2HDM_A (B) High-confidence predictions from Cfold generated ensembles with each structure’s average plDDT score (top), RMSD to the alternative crystal structure (middle), and RMSD to the dominant crystal structure (bottom). The color scheme of these structures matches the colorbar shown in panel A. Each row shows predicted alternative and dominant structures consistent with experiment and one high-confidence structure that is inconsistent. The experimentally determined gray alternative structure of Selecase was not predicted.

**Figure 3.**
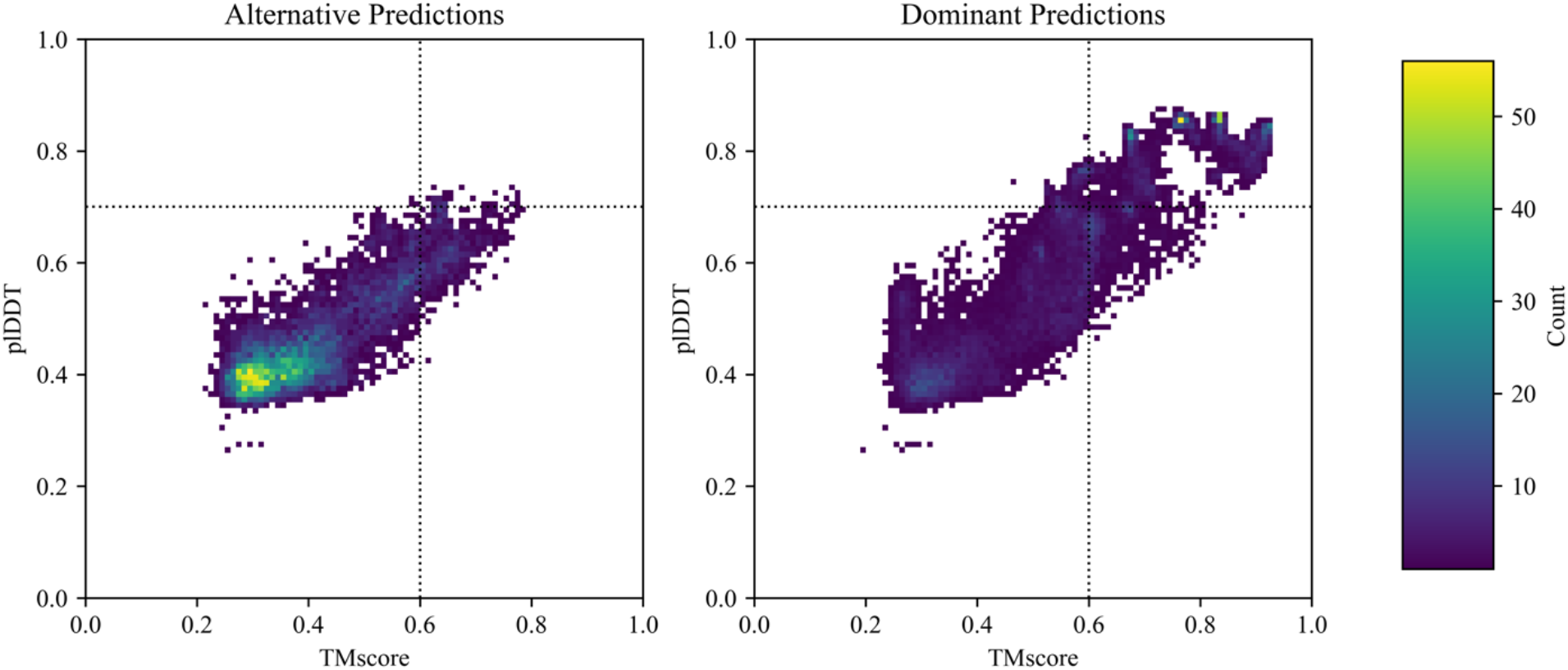
Two-dimensional histograms of all predicted structures from *Synthetic* GB98-T25I, *S. elongatus* KaiB, *R. sphaeroides* KaiB, *T. elongatus* KaiB, *H. sapiens* Mad2, *E. coli* RfaH, and *M. Jannashii* Selecase ensembles show distributions of plddt scores compared with TMscores. Predicted alternate structures appear on the left and dominant structures appear on the right (dominant/alternate defined in Figure). The colorbar corresponds to the number of predicted structures within a bin. Successful predictions have TM-scores ≥ 0.6 and plDDT scores ≥ 0.7; dashed lines are included as a visual aid. By these criteria, 27.9%/0.6% of dominant/alternative predictions successful.

While subfamily filtering enabled some successful predictions of alternative conformations, it also led to experimentally inconsistent predictions with high confidence. For instance, both experimentally consistent predictions of GB98-T25I in **Figure 2B** flank an experimentally inconsistent structure predicted with a confidence (plDDT = 0.82) substantially higher than the experimentally consistent helical structure (plDDT = 0.70). This type of misprediction is particularly difficult to recognize because the two-dimensional contact maps of the two predictions are nearly superimposable (**Figure S4A**). Thus, it is not obvious how coevolution could be used to discriminate between these two predictions, a situation that also occurs when using AlphaFold3 (Chakravarty et al. 2024). Similarly, CFold produces an experimentally inconsistent structure of KaiB with higher confidence than the experimentally consistent alternative structure (plDDT values of 0.7 and 0.72, respectively). Again, the two-dimensional representations of these conformations were superimposable, rendering coevolution unable to differentiate between the two predictions (**Figure S4B**). AF2 does not make either of these two mispredictions.

To determine whether CFold could distinguish between single folders and fold switchers, single-folding homologs of GB98-T25I, KaiB, and RfaH were also studied (**Figures S5, S6**). CFold incorrectly predicted that GA98, GB98, and GB98-T25I,L20A assume both the 3-α-helix bundle and an 4 β+α fold. However, these sequences are not known to switch folds, though their sequences are highly identical to GB98-T25I, which does. These findings show that AF2’s architecture can incorrectly misclassify singles folders as fold switchers with high confidence when their sequences are highly similar. CFold successfully predicted single-folding KaiBs and NusG as single folders, however, indicating that AF2’s architecture can distinguish between fold switchers and single folders with more disparate sequences.

Unlike AF2, which has been found to correctly predict fold switching with high confidence for all targets tested here except Selecase (Chakravarty et al. 2024), CFold predicts fold switching with high confidence for only GB98-T25I and one KaiB variant while also producing high-confidence incorrect predictions. Differences in training set best explain these predictive discrepancies since AF2 and CFold have the same network architecture, and their makers used very similar training approaches (Bryant and Noe 2024). The most direct explanation for AF2’s unique, high-accuracy predictions of these alternative conformations is that it recalls their structures from training (Bryant and Noe 2024; Chakravarty et al. 2024). An alternative explanation is that AF2’s more diverse training set helped it learn broader predictive principles than CFold’s smaller set allows. Although this explanation is possible, previous research shows that AF2 performs nearly as well with a training set of 10,000 structures as it does with its full set of over 100,000 (Ahdritz et al. 2024). Given that CFold was trained on >50,000 structures, it’s unclear if a larger dataset would yield additional insights. The most solid test would be to retrain AF2 on all conformations except for the alternative conformations presented here, but that requires very substantial computational resources.

CFold’s limited ability to successfully predict alternative structures outside of its training set supports the Associative Explanation and suggests several directions that may enable better predictions of yet-to-be-discovered alternative conformations in the future. First, since plDDT does not effectively discriminate between experimentally consistent and inconsistent structures predicted by either AF2 or CFold (Bryant and Noe 2024; Chakravarty et al. 2024), alternative measures need to be developed. Physically-based simulations and networks for reweighing members of predicted ensembles are promising avenues (Vani et al. 2024). Second, since subfamily filtering was the only approach that enabled predictions of alternative conformations tested here, methods to enhance that information may enable more accurate predictions of fold switching (Schafer and Porter 2023). Given previous observations, we were unsure of whether Finally, the problem of degenerate structural solutions to contact maps must be overcome. Since coevolution cannot always be used to discriminate between experimentally consistent and inconsistent predictions, incorporating physically-based priors into the predictive network may aid that effort (Ishizone et al. 2024). Data and code supporting this work can be found at: https://github.com/porterll/CFold_AF2.

## ACKNOWLEDGEMENTS

We thank Patrick Bryant for sharing the list of PDBs in CFold’s training set and Devlina Chakravarty for her feedback about this manuscript. This work utilized resources from the NIH HPS Biowulf cluster (http://hpc.nih.gov) and it was supported by the Intramural Research Program of the National Library of Medicine, National Institutes of Health (LM202011, L.L.P.).

