## Supplementary methods, tables and figures for "AlphaFold2’s training set powers its predictions of fold-switched conformations"

### Material and Methods

*Structure Prediction with CFold*

CFold was run with three different settings (Bryant and Noé 2024). The first set of predicted structures were generated using default settings and the num_samples_per_cluster flag set to 100. The second set of predicted structures were generated from multiple sequence alignment (MSA) sampling at each of the depths recommended by the authors of CFold: 16, 32, 64, 128, 256, 512, 1024 (Bryant and Noé 2024); 100 models were generated at each sampling depth. The final set of predictions take as input subfamily alignments from the original MSA and filtered by sequence identity and the num_samples_per_cluster flag set to 100. This is the same procedure followed by the ACE (Schafer and Porter 2023) pipeline to iteratively create subfamilies which contain sequences that are similar to the query sequence. To be thorough in our comparison we also use clustered MSAs (Wayment-Steele et al. 2024); however, CFold does not produce well folded alternative structures from these clusters (**Figure S2**). We generated 100 structures from each of the following sequence clusters from the AF-cluster GitHub repository: 1S2H_047, RFAH_049, WP_011056401.1, WP_027265801.1, WP_011056333.1, WP_069333443.1, WP_011242647.1. TM-scores were calculated with TM-align (Zhang and Skolnick 2005), and all-heavy-atom RMSDs with PyMOL with no outlier rejection (The PyMOL molecular graphics system, version 2.0 Schrödinger, LLC.). All structures were visualized with PyMOL.

*Determination of structures in CFold's training set*

We generated a FoldSeek database using a list of PDBs used to train CFold (Bryant and Noé 2024). We then used FoldSeek (van Kempen et al. 2024) to search this database for the structures of both conformations of all 8 proteins. Hits from searches, reported in **Table S1**, were considered valid if they had high FoldSeek scores (>$10^{-5}$), moderate TM-scores (0.6), and/or heavy-atom RMSDs ≥ 3.5Å.

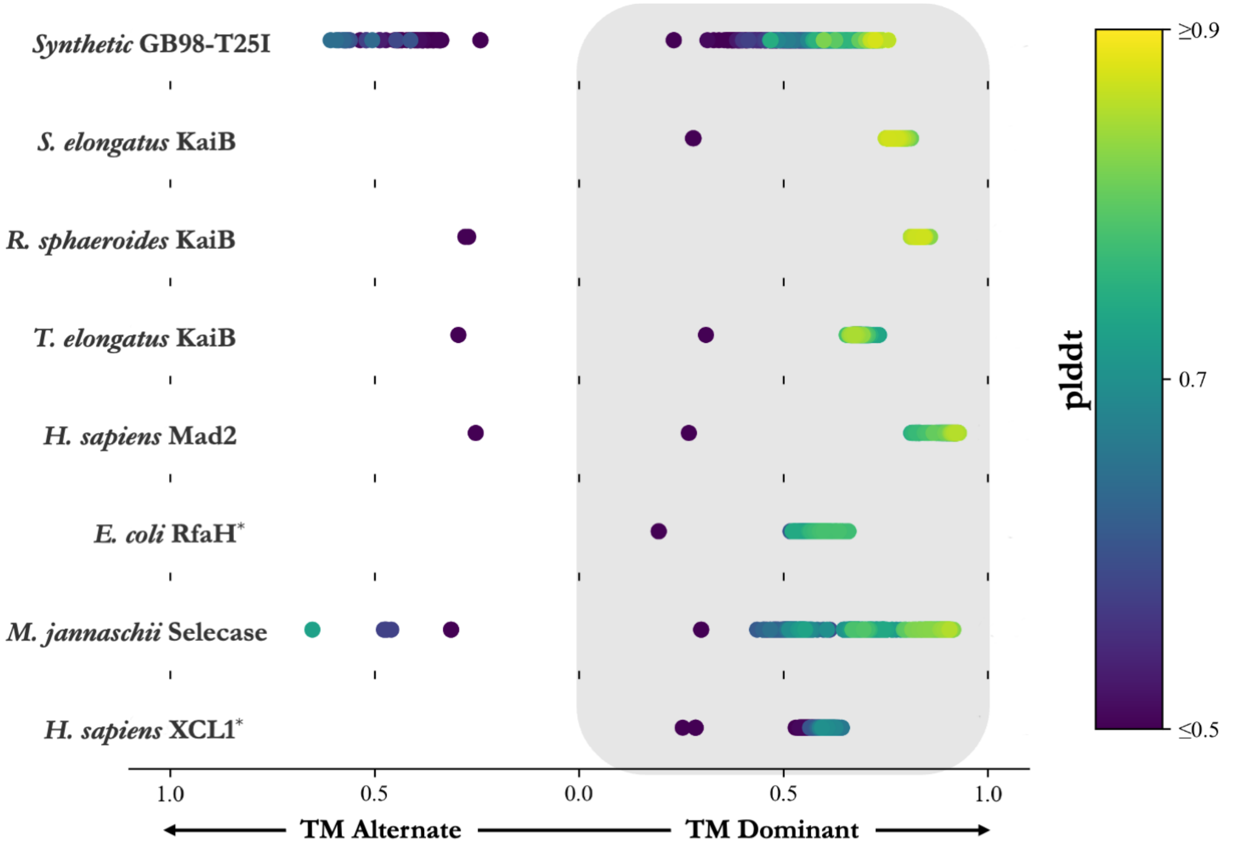


**Figure S1. TMscores for ensembles of predicted structures created with CFold for fold-switching proteins excluding predictions made with the subfamily alignments.** The predicted structure with the highest plddt score defines the dominant predicted conformation for the Cfold ensemble. Predicted structures within an ensemble are sorted by TMscore: if the TMscore is greater for the dominant conformation the value appears on the right side of panel A (gray box) and if the TMscore is greater for the alternative conformation the value appears on the left-side of panel A. All TMscores are colored by the predicted structures average plddt score. (* RfaH calculations are sorted by TMscore calculated for residues 118-155 to ensure conformations for this two-domain protein are organized by the fold-switching region. The TMscore plotted is the TMscore for the entire sequence compared to crystal structures. XCL1 calculations are on residues 1-65, the last 29 residues are unstructured and are not considered for TMscore or plddt score) Crystal structures used for comparison in order from top to bottom (alternative/dominant) are 2LHC/2LHD, 2QKE_A/1T4Y, 4KSO_A/8FWJ_M, 2QKE_A/1T4Y, 3GMH_L/2VFX_L, 5OND_A/6C6S_D, 4QHH_A/4QHF_A, 2N54_A/2HDM_A.

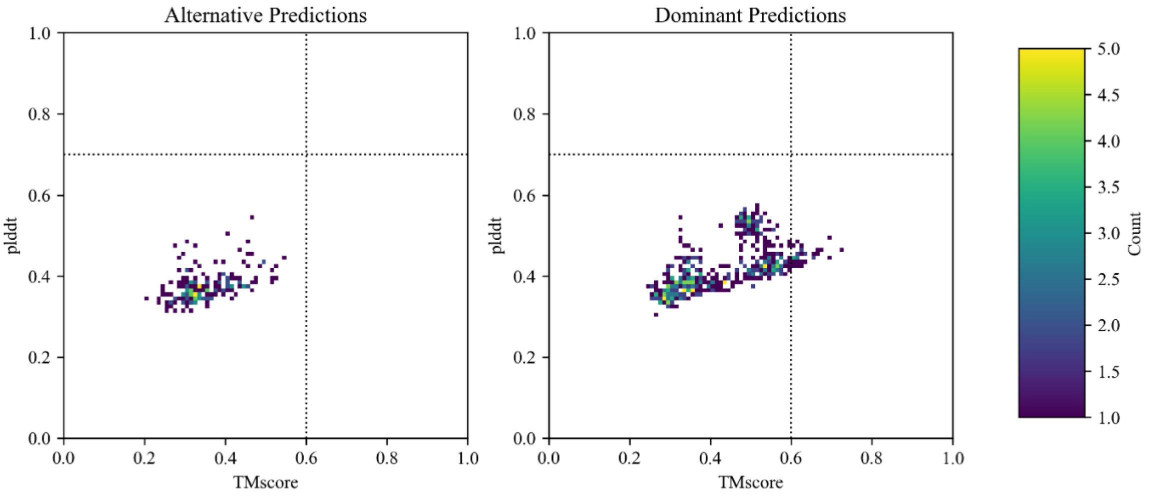


**Figure S2. 2D-Histogram of plddt scores versus TMscores for Cfold structures predicted using clustered MSAs.** Two-dimensional histograms of all predicted structures generated with AF-Cluster clusters from H. sapiens Mad2, L. pneumophila KaiB, R. sphaeroides KaiB, S. elongatus KaiB, T. elongatus KaiB, T. elongatus vestitus KaiB, E. coli RfaH ensembles show distributions of plddt scores compared with TMscores. Predicted structures within an ensemble are sorted by TMscore: if the TMscore is greater for the dominant conformation the value appears on the right side and if the TMscore is greater for the alternative conformation the value appears on the left-side. The colorbar corresponds to the number of predicted structures within a bin. A dashed line at TM-score = 0.6 and at plDDT = 0.7 are included as a visual aid.

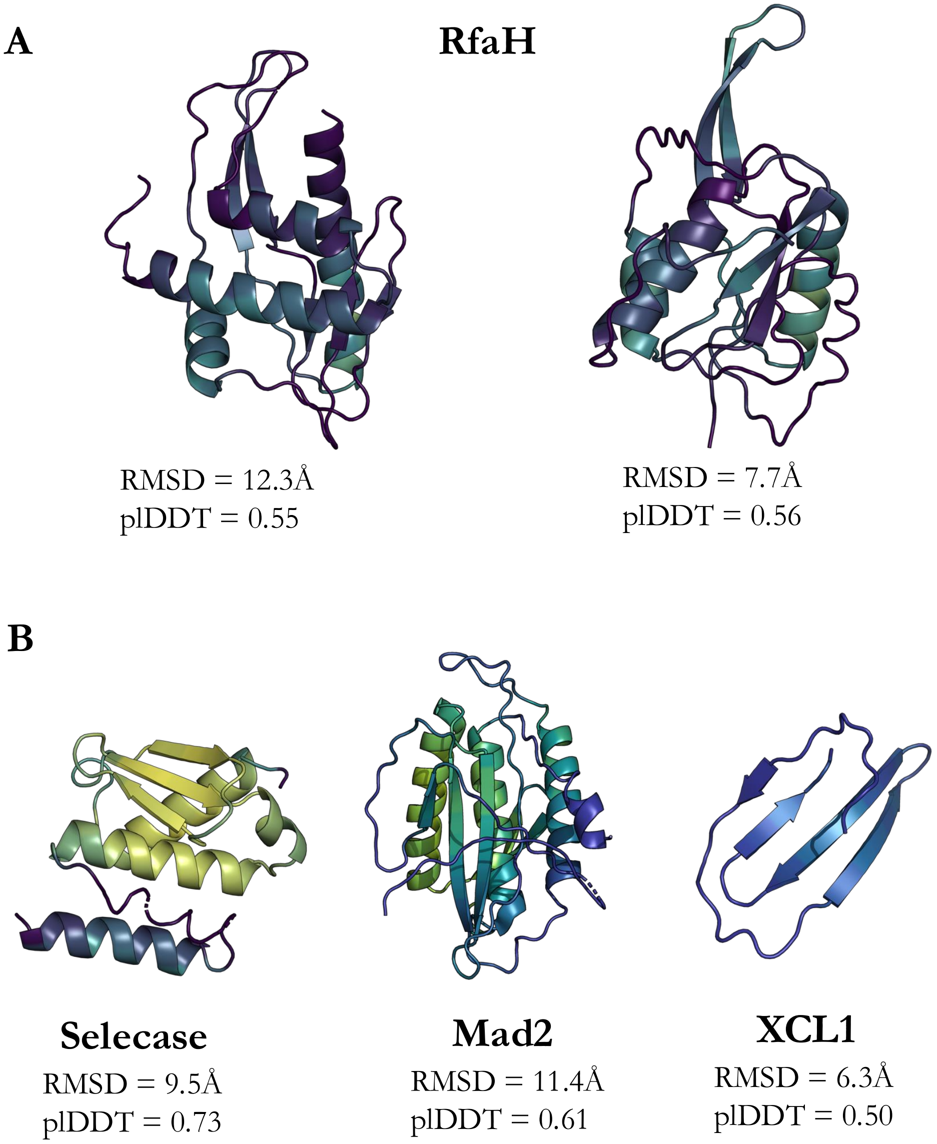


**Figure S3. CFold's best predictions of alternative fold-switched conformations are generally inaccurate and low-confidence. (A).** Sequence pruning enables CFold to predict an RfaH structure with a helical C-terminal domain (left), but its overall structure is inaccurate (>12.0 Å RMSD from experiment), and its confidence is low. Further, CFold produces an RfaH conformation with an experimentally inconsistent hybrid α-helix/β-sheet domain with equivalent confidence (right). **(B).** Despite extensive sampling of >2000 structures/protein, CFold predicts inaccurate alternative and low confidence conformations of Selecase, Mad2, and XCL1 at best. All models selected had the highest TM-scores to their respective experimentally determined alternative conformations. All-heavy-atom RMSDs are also calculated with respect to experimentally determined alternative conformations.

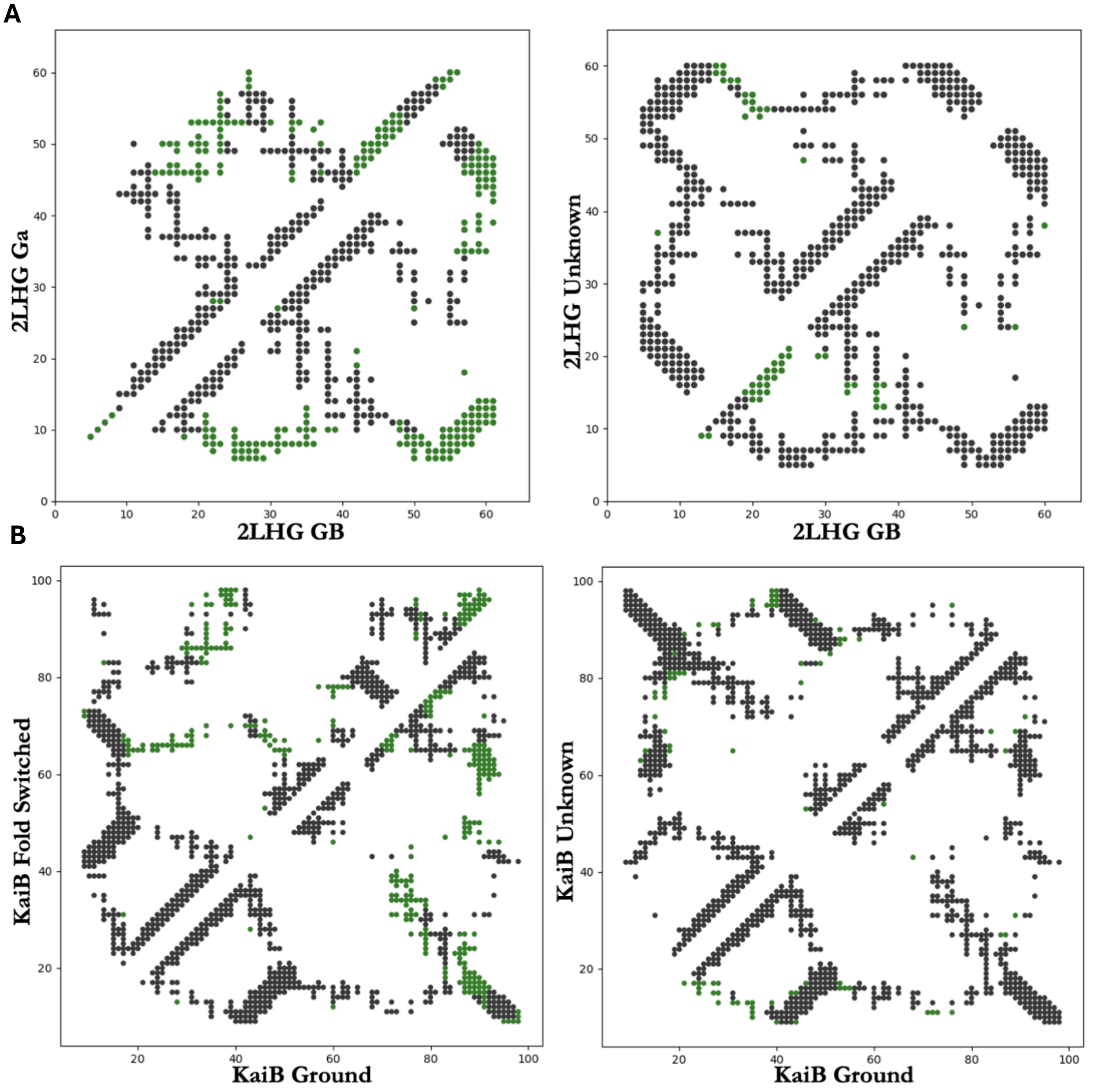


**Figure S4. Contact maps of experimentally consistent and inconsistent CFold predictions are indistinguishiable. (A)** Dual-fold contact maps for 2LHG structures predicted with Cfold. The dual fold contact map on the left-hand side shows the intra-molecular contacts for the GA conformation (upper triangle) and GB conformations' (lower triangle) residues with heavy atoms within 8Å. Black Contacts are common between the two conformations and green contacts are unique to that conformation. The dual fold contact map on the right-hand side shows the intra-molecular contacts for the GB conformation (lower triangle) compared to the experimentally inconsistent alternative conformation. Despite the three-dimensional conformations being distinct the two-dimensional representations are nearly degenerate. **(B)** Dual-fold contact maps for *R. sphaeroides* KaiB structures predicted with Cfold. The dual fold contact map on the left-hand side shows the intra-molecular contacts for the fold-switched (upper triangle) and ground state conformations' (lower triangle) residues with heavy atoms within 8Å. Black Contacts are common between the two conformations and green contacts are unique to that conformation. The dual fold contact map on the right-hand side shows the intra-molecular contacts for the GB fold (lower triangle) compared to the experimentally inconsistent alternative conformation. Despite the three-dimensional conformations being distinct the two-dimensional representations are nearly degenerate.

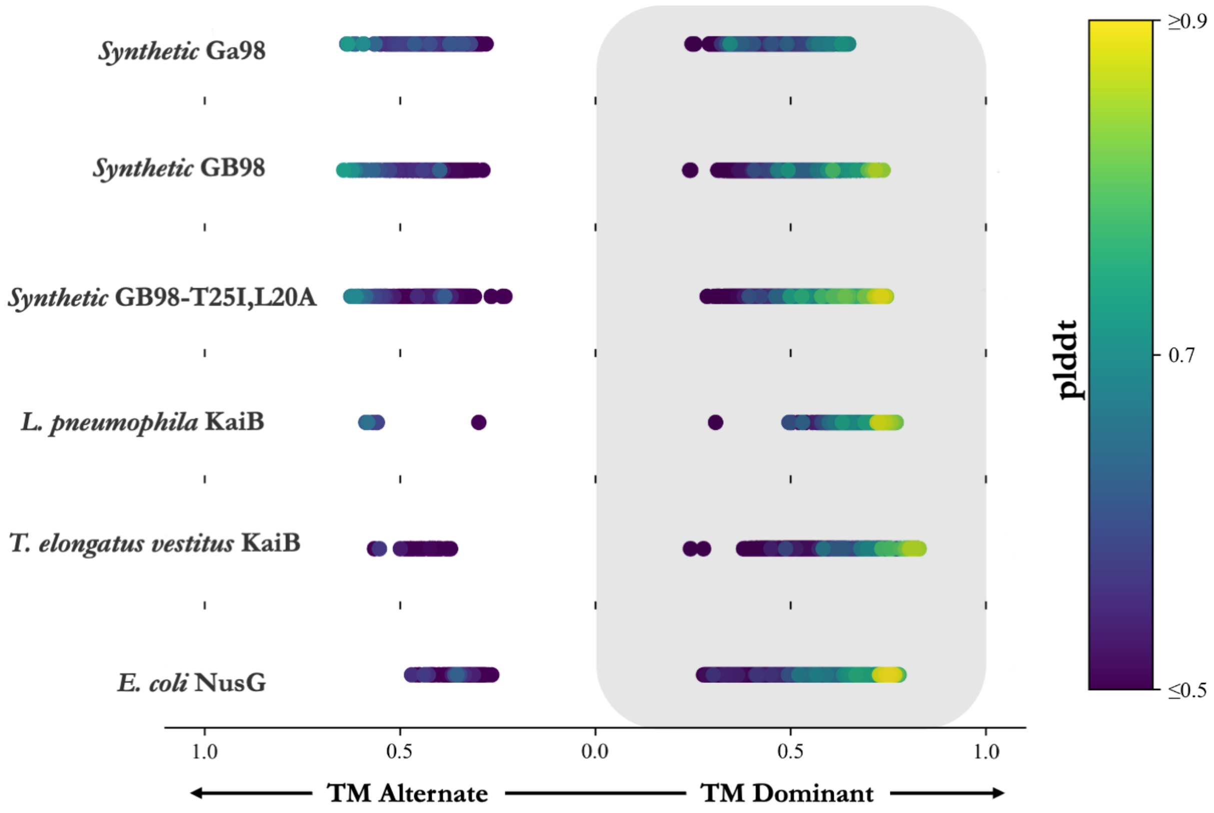


**Figure S5. TMscores for ensembles of predicted structures created with Cfold for single-folding homologs of fold-switching proteins.** The predicted structure with the highest plDDT score defines the dominant predicted conformation for the CFold ensemble. Predicted structures within an ensemble are sorted by TMscore: if the TMscore is greater for the dominant conformation the value appears on the right side (gray box) and if the TMscore is greater for the alternative conformation the value appears on the left side. All TMscores are colored by the predicted structures average plddt score. Crystal structures used for comparison in order from top to bottom (alternative/dominant) are 2LHD/2LHC, 2LHC/2LHD, 2LHC/2LHD, 2QKE_A/1T4Y, 2QKE_A/1T4Y, 5OND_A/6C6S_D

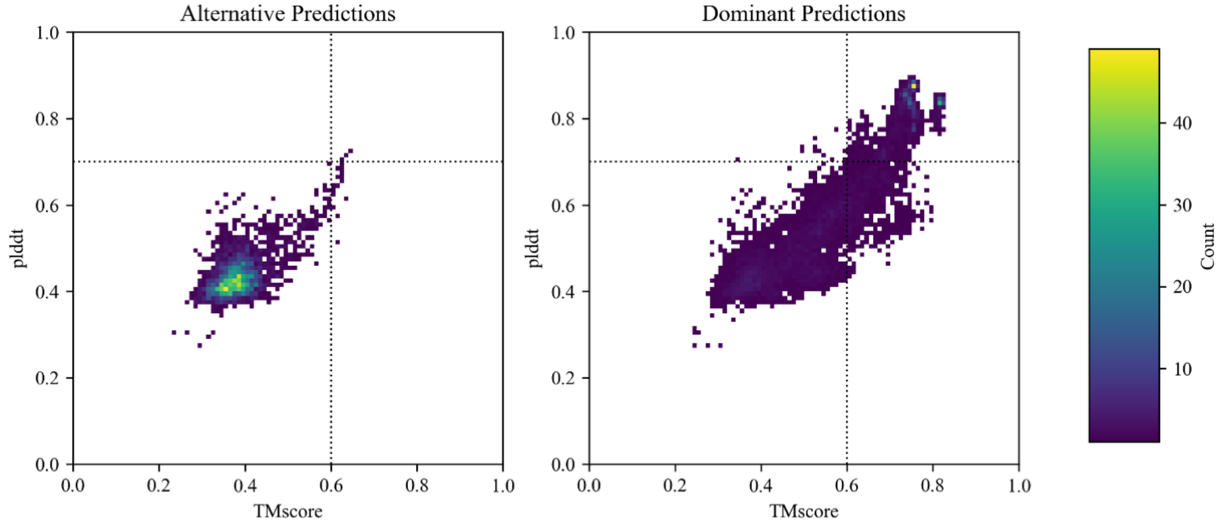


**Figure S6 2D-Histogram of plddt scores versus TMscores for Cfold structures.** Two-dimensional histograms of all predicted structures from Synthetic GA98, Synthetic GB98, Synthetic GB98-T25I, L20A, L. pneumophila KaiB, T. elongatus vestitus KaiB, E. coli NusG ensembles show distributions of plDDT scores compared with TMscores. Predicted structures within an ensemble are sorted by TMscore: if the TMscore is greater for the dominant conformation the value appears on the right side and if the TMscore is greater for the alternative conformation the value appears on the left-side. The colorbar corresponds to the number of predicted structures within a bin. A dashed line at TM-score = 0.6 and at plddt = 0.7 are included as a visual aid.

Table S1. Protein conformations represented in CFold's training set. Unlisted conformations were not found in the training set.

| Protein | Training set structure |
|---|---|
| GA | 4HEO |
| GB | 6L9B |
| KaiB (dominant) | 4HU7 |
| Mad2 (dominant) | 6KEA |
| RfaH (dominant) | 2XHC |
| Selecase (dominant) | 6SAR |
| XCL1 (dominant) | 1B3A |

Table S2. Number of structures produced for each protein sequence with the Cfold algorithm, Cfold run with pruned subfamily MSAs as input, and a single sequence as input.

| ID | Cfold | Subfamilies | Single sequence |
|---|---|---|---|
| ***Synthetic* GB98-T25I** | 800 | 1900 | 4 |
| ***Synthetic* Ga98** | 800 | 1100 | 4 |
| ***Synthetic* GB98** | 800 | 1400 | 5 |
| ***Synthetic* GB98-T25I,L20A** | 800 | 1400 | 5 |
| ***S. elongatus* KaiB** | 800 | 1800 | 3 |
| ***R. sphaeroides* KaiB** | 800 | 1100 | 2 |
| ***T. elongatus* KaiB** | 800 | 2000 | 2 |
| ***L. pneumophila* KaiB** | 800 | 400 | 2 |
| ***T. elongatus vestitus* KaiB** | 800 | 1100 | 3 |
| ***E. coli* RfaH** | 800 | 3000 | 2 |
| ***E. coli* NusG** | 800 | 1300 | 3 |
| ***M. jannaschii* Selecase** | 800 | 1200 | 4 |
| ***H. sapiens* XCL1*** | 800 | 3400 | 4 |
| ***H. sapiens* Mad2** | 800 | 2800 | 2 |
